# Beyond completion time: predicting individual differences in executive functions using cursor trajectory features in an online Trail Making Test

**DOI:** 10.64898/2026.08.04.742901

**Authors:** Gustavo E. Juantorena, Gianluca Capelo, Juan E. Kamienkowski

## Abstract

**Background:** The Trail Making Test (TMT) is a widely used instrument for assessing executive functions due to its sensitivity. But, its traditional scoring, based solely on total completion time, limits its specificity by losing the rich behaviour required to complete the task, involving integrating visual search, motor planning and task switching. A computerised implementation of the TMT (cTMT) allows high-resolution cursor trajectories to be recorded, offering access to fine-grained features of the trajectory, while an online administration improves accessibility and statistical power.

**Objective:** This study aimed to extract high-resolution cursor trajectory features from an online cTMT and evaluate their capacity to predict individual differences in core executive functions, such as visual working memory (VWM), inhibitory control and age, using machine learning, as well as validating the feasibility of a fully online data acquisition pipeline as a basis for digital biomarkers.

**Methods:** Participants completed an online battery comprising the cTMT and three validation tasks: the Change Detection Task (CDT), the Stop-Signal Task (SST) and the Go/No-Go task (GNG). A total of 104 features (26 features x Part A/B x whole-trial/first-10-targets) were extracted from cursor trajectories, including trajectory profiles and segmentation into latent states: Search, Travel and Hesitation. Regression models were trained within a nested Leave-One-Out cross-validation framework with inner 10-fold hyperparameter tuning, feature selection and standardisation applied strictly within folds. Performance was assessed via mean absolute error (MAE), normalised error (MAE/SD) and permutation testing; SHapley Additive exPlanations (SHAP) were used to characterise feature importance.

**Results:** cTMT features strongly predicted age across all models (p < .001), with MAEs roughly one-third lower than the target’s dispersion, driven predominantly by distance-based “circuitousness” metrics from both parts. GNG accuracy and c-coefficient were both significantly predicted, but with a dissociated pattern: accuracy was best explained by Part B (alternation) metrics, whereas the c-coefficient was almost exclusively predicted by Part A (simple sequencing) metrics. SST response time (SSRT) was not significantly predicted by any model. VWM, characterised by the mean Cowan s K, was significantly predicted mainly by the more complex models, and involving state transitions and search phases in both Part A and B.

**Conclusions:** Moving beyond completion time, cursor trajectory dynamics from an online cTMT provide a rich behavioural signal for predicting age, VWM capacity and distinct aspects of inhibitory control. The dissociation between Part A and Part B predictors supports differentiated cognitive processes within the TMT and positions the cTMT as a scalable, portable digital biomarker with promise for computational psychiatry and personalised neuropsychology.

## I. Introduction

The emergence of digital neuropsychology ^1^ and computational psychiatry ^2^ has enabled new ways to assess cognition using online platforms and high-resolution behavioral measurements. Unlike traditional in-person or paper-based evaluations, these approaches employ standard digital devices to deliver scalable, cost-efficient tools capable of recording high-resolution behavioral signals. Digital methods not only improve accessibility but also make it possible to extract latent cognitive variables with greater precision by analyzing continuous data streams, such as reaction times, trajectory dynamics, and precise motor patterns ^3,4^. and machine learning techniques to identify subtle behavioral signatures linked to cognitive processes.

Within this context, executive functions are a core domain of cognitive science and are essential for goal-directed behavior. These functions include inhibitory control, cognitive flexibility, and working memory ^5,6^. Operating in a top-down manner, these processes allow individuals to override habitual responses, maintain task goals, and coordinate actions involving multiple steps. Executive functions are critical for everyday activities, such as planning, decision-making, and adaptive behavior. Deficits in these abilities often appear early in neurodevelopmental and neurodegenerative conditions, making them valuable targets for early detection.

A standard instrument for assessing these abilities is the Trail Making Test (TMT). It has wide use in clinical and research settings for the evaluation of executive functions. It consists of two parts. In Part A, participants connect numbers in ascending order. In Part B, participants alternate between connecting numbers and letters. While the test is highly sensitive to executive dysfunction, its traditional scoring system, which is based solely on total completion time, limits its specificity ^7–11^.

However, this reliance on a single completion-time measure highlights an important limitation of the traditional format. The richness of the behavior involved in the TMT, which integrates visual search, motor planning, hand control, and task switching, is largely lost when only a single global metric is retained. The computerized implementation of the TMT (cTMT) provides a solution by enabling the analysis of several fine-grained metrics derived from precise motor and cursor movements, offering access to features such as pauses, trajectory deviations, speed fluctuations, and search patterns.

In addition to these analytical advantages, the possibility of online experimentation offers practical benefits for data collection. On the one hand, diverse samples can be rapidly collected in natural environments, improving the ecological validity and the accessibility of populations other than WEIRD ^12^. On the other hand, it increases statistical power. The digital version of the TMT was released online, as well as complementary tasks that measure working memory and inhibitory control. This allows us to investigate whether features extracted from cTMT cursor trajectories can predict variations in core executive functions. Understanding these relationships can improve theoretical models of cognitive control, and contribute to the development of scalable digital biomarkers for the early detection of executive dysfunction in the general population.

The digitization of the Trail Making Test (TMT) has expanded its clinical utility across populations and conditions. Digital versions sensitively detect age-related cognitive differences ^13^ and support early diagnosis of Alzheimer’s ^14^ and Parkinson’s disease ^15^. Advanced analytical approaches, including hidden Markov models, extract richer cognitive markers beyond completion time ^16^, while mobile and tablet-based implementations have demonstrated ecological validity ^17,18^. Early foundational work ^19^ and recent findings on set-size effects ^20^ further inform the cognitive mechanisms captured by the task.

To rigorously evaluate this predictive potential, we included a battery of standardized tasks to serve as ground truth measures for specific executive domains. We assessed Visual Working Memory (VWM) capacity using the Change Detection Task ^21^. For inhibitory control, we employed two distinct paradigms to capture different facets of inhibition: the Go/No-Go task ^22,23^, targeting action restraint (withholding a response), and the Stop-Signal Task ^24^, designed to measure reactive inhibition (cancelling an ongoing action). Performance on these established benchmarks serves as the target variable for the predictive models.

The present research is driven by two complementary goals. From a theoretical perspective, we aim to understand the cognitive mechanisms underlying complex task execution through high-precision measurements of motor output. From an applied perspective, we seek to translate this understanding into a sensitive, precise, and portable tool for the assessment of executive deficits.

To achieve these general goals, the present study focuses on three specific objectives: first, we aim to extract features from cursor trajectories, such as hesitation periods and velocity profiles, which are not captured in traditional assessments. Second, we seek to evaluate the ability of these metrics to predict visual working memory capacity, inhibitory control, and age using machine learning techniques. Finally, we aim to validate the feasibility of the data acquisition pipeline in an online environment and contribute to the development of digital biomarkers.

## II. Methods

### A. Participants

A total of 484 unique participants were identified across the four cognitive tasks (cTMT, SST, CDT, and Go/No-Go) after applying task-specific data quality controls. Not all participants completed every task: 386 completed the cTMT, 246 the SST, 248 the CDT, and 310 the Go/No-Go task. Participants were recruited at the university and through social media. All participants were naive to the experiment’s objectives, older than 18 years, and willingly provided electronic informed consent. Ethical approval for the study was obtained from the respective Ethics Committees of each university Protocol 295 from the Instituto de Investigaciones Médicas “Alfredo Lanari” (University of Buenos Aires).

Of the 484 participants that completed at least one task, self-reported demographic metadata was collected (Fig. 1). Gender was available for 471 participants, age was available for 457, nationality was available for 471, and education was available for 462 participants. The cohort exhibited a balanced gender distribution, with a slight prevalence of female participants (N=234, 49.7%) compared to males (N=226, 48.0%); a minority identified as gender diverse (N=11, 2.3%). The age of participants ranged from 18 to 77 years [M=32.1, SD=10.4], with the sample consisting predominantly of young adults. Information regarding nationality, country of residence, and native language were collected. Regarding nationality, the sample was predominantly Argentinean (N=383, 81.3%), with smaller representations from other Latin American countries including Uruguay (N=23), Colombia (N=14), Venezuela (N=11), and Peru (N=10). In terms of education, the sample was highly educated: the largest group possessed a completed university degree (N=207, 44.8%), followed by those with completed secondary education (N=157, 34.1%). Smaller subsets reported having completed postgraduate degrees (N=56, 12.2%), tertiary/vocational training (N=36, 7.8%) or primary education only (N=6, 1.3%).

**Fig. 1.**
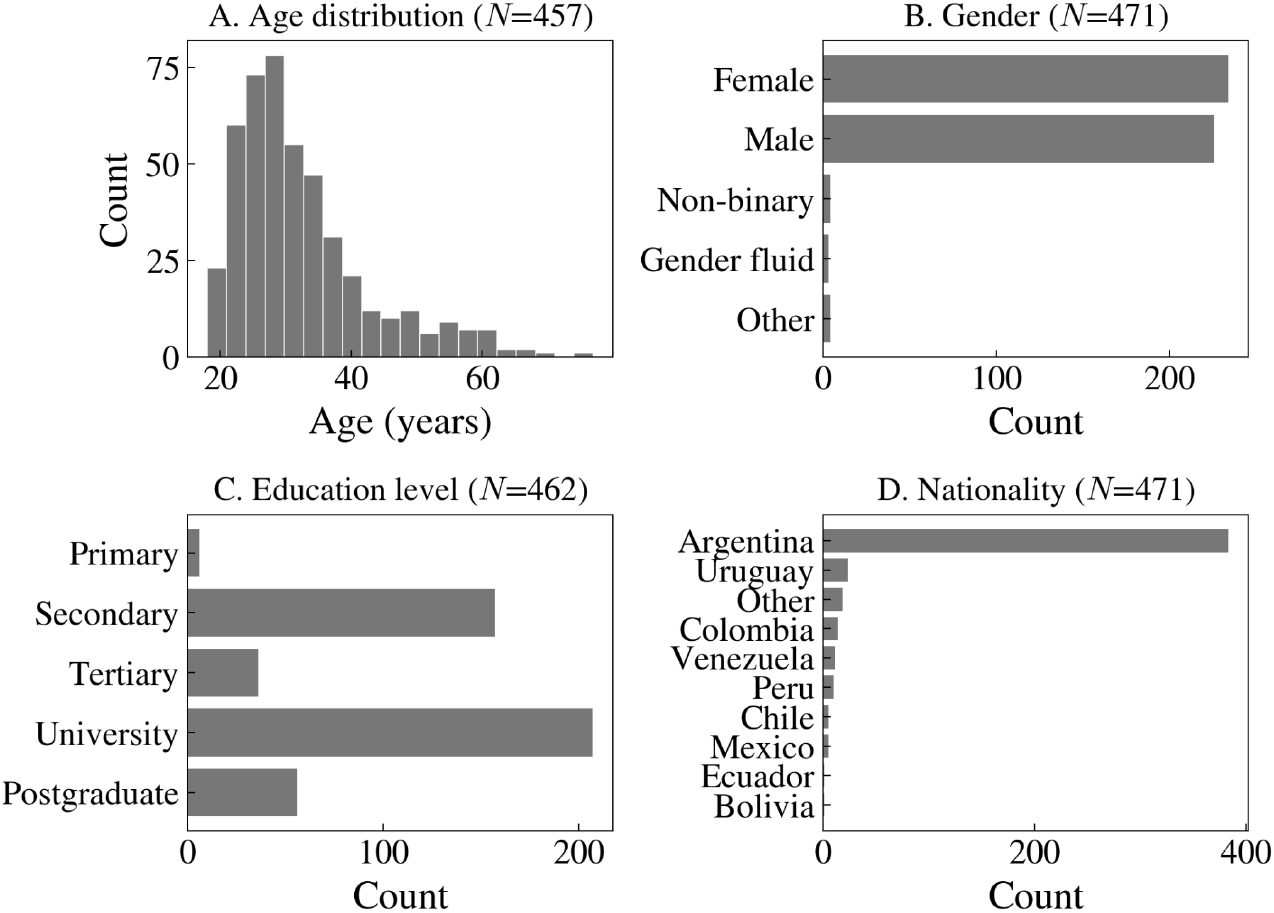
**(A)** Age Distribution of Participants Exhibiting a Right-Skewed Profile Prevalent in Young Adulthood. **(B)** Highest Educational Attainment Levels **(C)** Geographic distribution by country of origin. **(D)** Self-reported gender identity. Variations in sample sizes across panels (457–471) reflect the availability of valid data after quality control filtering. The data shown represent valid responses for each category.

### B. Inclusion criteria

The study only included subjects with no history of psychiatric disorders, epilepsy, seizures, drug or alcohol abuse, or psychotropic medication use. Additionally, they were asked whether they have normal or corrected-to-normal vision. Furthermore, only individuals aged 18 or older were admitted. Finally, the experiment could not be completed on mobile phones, tablets, or screens smaller than 13 inches.

### C. Web platform and implementation details

Participants accessed the study through DataPruebas (https://datapruebas.org/), a Spanish-language web platform ^25^. This platform allows researchers to set experiment parameters such as duration, number of allowed attempts, technical requirements (e.g., webcam availability, device type, or minimum screen size), and eligibility criteria (e.g., age, location), while enabling real-time monitoring of participant progress via an execution dashboard. Upon entering the platform, participants provided informed consent and completed a demographic questionnaire. They were then instructed to complete the four cognitive tasks described below, which could be performed across separate sessions.

We developed the tasks using jsPsych (v6.3; ^26^), a JavaScript library designed for creating online behavioral experiments. The source code for all tasks is available at https://github.com/NeuroLIAA/Neuro_psych_tasks. To ensure consistent stimulus dimensions across participants using different monitors, we employed the jspsych-virtual-chinrest plugin, available in the jsPsych library (v6.3.0+). The procedure consisted of two steps designed to estimate the viewing distance and standardize the screen resolution. First, participants were presented with an on-screen rectangle and instructed to adjust its size to match a standard physical card (e.g., a credit card conforming to the ID-1 standard, approx. 85.60 × 53.98 mm). This step allowed us to calculate the pixel density of the user’s display, relating digital pixels to physical measurements ^27^. Subsequently, a blind spot detection task estimated viewing distance (3 repetitions, averaged). Using these two measures and trigonometric principles, stimuli were dynamically resized to maintain constant physical dimensions across participants.

### D. Experimental Tasks

#### Trail-Making Test

We implemented an online version of the Trail Making Test (cTMT) using the jsPsych library and a custom-built plugin (Fig. 2A). Participants performed two types of trials: cTMT-A, requiring them to connect numbers (1-20) sequentially, and cTMT-B, requiring alternating connections between numbers and letters (1-A-2-B…). Each trial type was primed by a distinct color cue (blue fixation cross for cTMT-A, red for cTMT-B). The protocol included a training phase (one trial per condition) where targets were visually highlighted upon interaction to familiarize participants with the mechanics, followed by a testing phase consisting of 10 trials per condition without visual feedback. Participants were instructed to use a mouse to complete the trails as quickly and accurately as possible. In the event of an error, participants were required to return to the last valid target to resume the path. The system recorded total completion time, accuracy, and high-resolution cursor trajectories (x, y coordinates and timestamps). Only trials from the testing phase were included in the analysis.

**Fig. 2.**
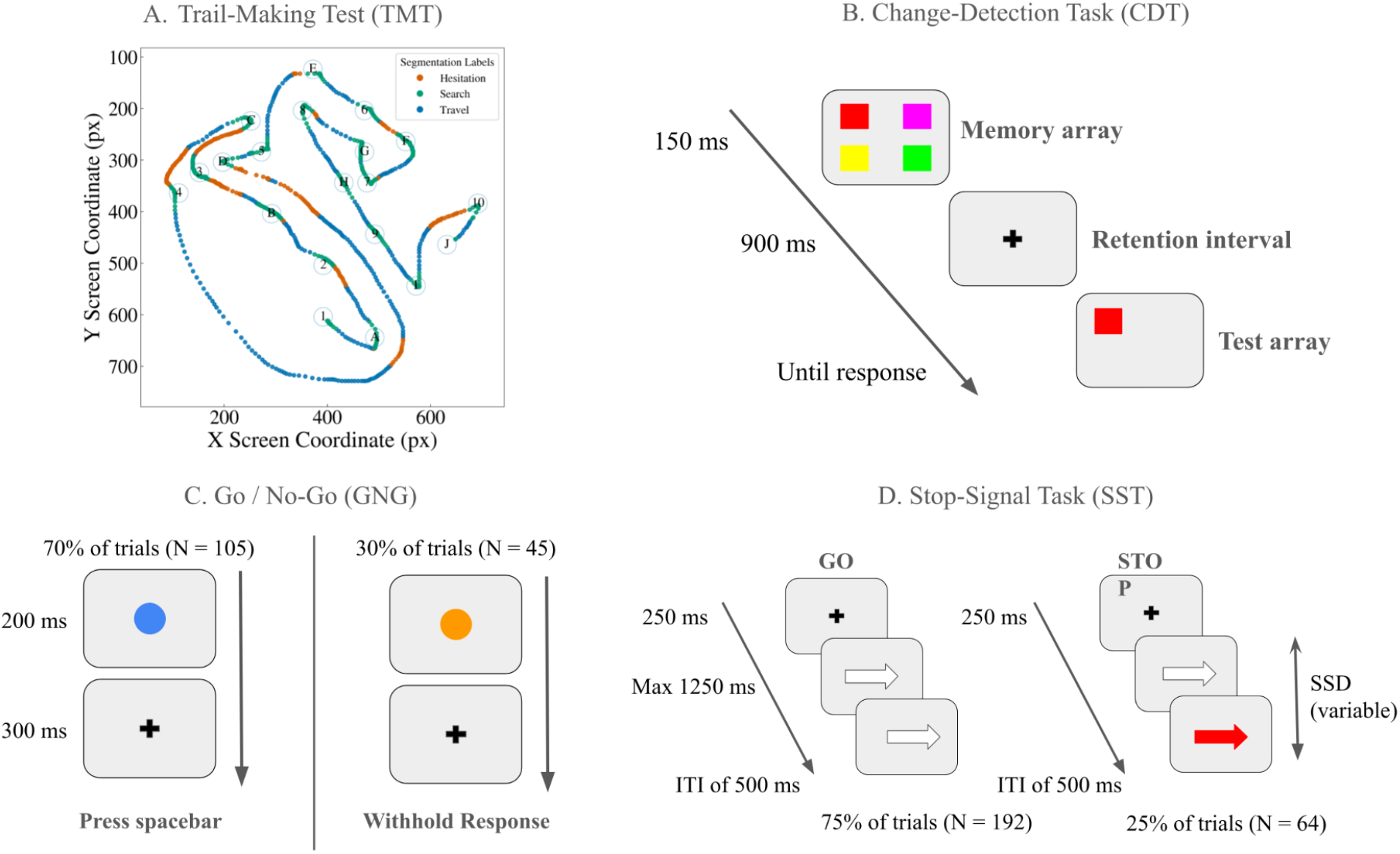
Graphical representation of the four cognitive tasks implemented on the platform. **(A)** Computerized Trail-Making Test (cTMT), showing the setup for tracking hand path and cursor trajectories. The numbered circles represent the targets. This example corresponds to cTMT-B, alternating numbers and letters in alphabetical order. The scatter plot shows the sequential path of the participant’s mouse between them. The color of the points indicates the state: hesitation (blue), search (orange), or travel (green) (see “Feature Extraction and Segmentation” section). **(B)** Change-Detection Task (CDT), evaluating visual working memory. **(C)** Go/No-Go task, assessing action restraint. **(D)** Stop-Signal Task (SST), measuring reactive inhibition.

#### Change-Detection Task

Visual working memory (VWM) capacity was assessed using a change detection paradigm implemented via a custom-built jsPsych plugin (Fig. 2B). The design followed the canonical protocol established by Luck and Vogel ^21^. Each trial began with a central fixation cross, followed by an encoding array of either 4 or 6 colored squares presented for 150 ms. The squares were drawn from a pool of eight highly discriminable colors (black, blue, cyan, green, magenta, orange, red, and yellow). Following a retention interval of 900 ms, a single probe item appeared at the location of one of the previous items.

Participants were required to judge whether the color of the probe matched the item originally presented at that specific location. Responses were registered via keyboard (using ‘L’ for “Same” and ‘D’ for “Different”) within a maximum window of 5000 ms. The session began with a training block with feedback, followed by a testing phase consisting of 120 trials divided into three blocks. Trials were balanced across set sizes (60 trials at set size 4; 60 at set size 6) and change conditions (50% match probability). The primary outcome measure was memory capacity (Cowan’s K) (Table 1), alongside accuracy and response time.

**Table 1.** Summary of the outcome measures used as prediction targets, derived from the three validation tasks and demographic data. For each target, the formula and cognitive interpretation are provided. HR: Hit Rate; FA: False Alarm rate; HR, FA: sample means; SDHR,S DFA: sample standard deviations; S: set size; SSD: Stop Signal Delay; pr: probability of responding on stop trials.

| Task | Target name | Formula | Interpretation |
| --- | --- | --- | --- |
| Change Detection Task | Cowan's K | $K = S \cdot (2 \cdot Accuracy - 1)$ | **Visual working memory capacity:** higher K enables more fluid navigation and efficient search patterns |
| Stop Signal Task | SSRT | $SSRT = F_{Go}^{-1}(p_r) - \overline{SSD}$ | **Motor inhibitory control:** longer times indicate greater difficulty suppressing a prepotent response. |
| Go/No-Go | $c$-Coefficient | $c = -0.5 \cdot \left( \frac{HR - \overline{HR}}{SD_{HR}} + \frac{FA - \overline{FA}}{SD_{FA}} \right)$ | **Response bias:** linked to baseline scanning efficiency and planning decisiveness in trajectory execution |
| Go/No-Go | Accuracy | $Accuracy = \frac{Correct}{N_{trials}}$ | **Overall performance under inhibitory challenge:** balancing 'Go' trial hits against 'No-Go' false alarms |
| Demography | Age | $Age [18, 77] \text{ y.o.}$ | **Brain aging:** characterized by increased motor noise, reduced spatial efficiency, and longer motor planning periods |

#### Go/No-Go

We implemented a Go/No-Go paradigm using jsPsych to assess action restraint ^28^, the ability to withhold a response before it is initiated (Fig. 2C) ^29^. Participants were presented with a series of colored circles in the center of the screen and were instructed to press the spacebar as quickly as possible when a blue circle appeared (“Go” condition) and to withhold their response when an orange circle appeared (“No-Go” condition).

The experiment began with a training block (6 trials) featuring slower timing and immediate feedback. The subsequent testing phase consisted of 150 trials divided into 15 blocks. Crucially, the probability of “Go” signals was set at 70% (105 trials) versus 30% for “No-Go” signals (45 trials); this high frequency of Go trials was designed to establish a prepotent tendency to respond, thereby increasing the cognitive demand on inhibitory restraint mechanisms.

During the test phase the circle was presented for 200 ms. Immediately following the offset of the stimulus, a fixation cross appeared for 300 ms, resulting in a total response window of 500 ms per trial. If a participant responded within this window or if the time elapsed, the trial concluded.

#### Stop-Signal Task

The Stop-Signal Task was employed to assess reactive inhibition, defined as the ability to cancel an ongoing motor response upon the presentation of an unexpected stop signal. This implementation was based on the STOP-IT code (^30^; https://github.com/fredvbrug/STOP-IT). The task is grounded in the independent race model ^24^, which conceptualizes performance as a race between two competing processes: a “Go” process triggered by the primary stimulus and a “Stop” process triggered by the stop signal (Fig. 2D).

Participants were instructed to respond as quickly and accurately as possible to a “Go” stimulus (a white arrow pointing left or right) by pressing the corresponding arrow key on the keyboard. On a minority of trials (25%), the “Go” stimulus changed color to red (the “Stop” signal) after a variable delay, indicating that the participant should inhibit their response. The task consisted of one practice block of 32 trials and four experimental blocks of 64 trials each, resulting in a total of 256 experimental trials. Each trial began with a fixation cross (250 ms), followed by the stimulus presentation (maximum duration 1250 ms) and a fixed inter-trial interval (ITI) of 500 ms. Crucially, participants were explicitly instructed not to delay their responses in anticipation of the stop signal, in order to maintain valid reaction time distributions for the “Go” process.

To ensure an approximately equal distribution of successful and failed inhibition trials (a prerequisite for the race model) the Stop Signal Delay (SSD) was adjusted dynamically using a 1-up/1-down adaptive staircase algorithm. The initial SSD was set to 200 ms and adjusted by ± 50 ms following successful and failed inhibition trials, respectively. Feedback on mean reaction time and accuracy was provided during the break periods between blocks to allow participants to monitor their performance.

### E. Data Processing and Feature Extraction

#### Data Quality Checks

To mitigate the risks associated with unsupervised online data collection and ensure the reliability of the behavioral measures, we implemented data quality checks. First, participants registered under a unique persistent identifier, preventing duplicate submissions and enabling data linkage across tasks. Second, to standardize the physical experimental conditions, we employed the Virtual Chinrest procedure. This allowed us to estimate viewing distance and screen size, enabling us to exclude participants with displays too small to render the cTMT spatial layout correctly. Furthermore, participation was restricted to desktop or laptop users; we explicitly excluded touchscreen devices to ensure that the recorded trajectories reflected precise mouse or trackpad dynamics rather than touch interactions. Third, we applied performance filters to remove low-effort or automated responses. For the cTMT, valid trials were defined as those where the participant successfully connected a minimum of 10 targets; trials falling below this threshold were excluded. A total of 386 participants completed the cTMT. Eighteen participants (4.7%) were excluded from subsequent analyses due to incomplete data: 16 participants lacked valid cTMT-B trials (Part B), and one lacked valid cTMT-A trials (Part A). The final sample comprised 368 participants with complete cTMT data (i.e., at least one valid trial for both Part A and Part B). Finally, for the reaction-time sensitive tasks (SST, GNG, CDT), we applied a physiological lower bound filter of 150 ms. While Whelan ^31^ notes that genuine reaction times typically have a physiological minimum of at least 100 ms, we selected a conservative threshold of 150 ms within his recommended 100–200 ms exclusion range to rigorously remove anticipatory responses.

#### cTMT Trajectory Preprocessing

Raw cursor time-series (x, y coordinates and timestamps) from the cTMT were processed using *Neurorask*, a custom-built Python library designed for high-resolution behavioral analysis https://github.com/NeuroLIAA/neurotask. To ensure data quality, trials with fewer than ten connected targets were excluded.

Because data was collected via web browsers, the raw sampling rate was inherently irregular due to variations in browser event firing, system load, and hardware performance. To standardize the temporal resolution across all trials and participants, we applied a linear interpolation procedure to the trajectories. First, duplicate timestamps were removed to ensure temporal monotonicity, and then, the *x* and *y* coordinates were then independently resampled onto a uniform 60 Hz temporal grid (approx 16.67 ms intervals). This frequency was selected to align with the standard refresh rate of most consumer displays and to provide a sufficient sampling rate to capture the bandwidth of intentional human motor output without introducing high-frequency noise. This procedure established a consistent temporal resolution while preserving the spatial and kinematic integrity of the original movement data. For incomplete but valid trials (≥10 targets), features were calculated based on the trajectory up to the last correctly connected node to ensure comparability across the sample.

#### Feature Extraction and Segmentation

We extracted a high-dimensional feature set categorized into three distinct domains (see Table S1 in supplementary materials). Regarding performance and temporal metrics, we calculated intra-target time (dwell time) and inter-target time (transition time). In terms of kinematics and dynamics, we derived velocity and acceleration profiles to compute mean and peak speeds. Furthermore, to quantify spatial error and motor control precision, we compared the participant’s actual path against an optimal path consisting of linear connections between consecutive targets. From this comparison, we extracted metrics including the cumulative point-to-point Euclidean deviation and the area under the curve (AUC) between the actual and optimal trajectories. To capture latent cognitive processes during task execution, we implemented a trajectory segmentation algorithm adapted from ^17^. The continuous cursor stream was discretized into three mutually exclusive states based on a participant-specific velocity threshold, defined as the median speed between the first two targets. These states include *Search*, characterized by low-velocity periods where the cursor is stationary or moving blindly while the user visually scans for the next target; *Travel*, consisting of high-velocity ballistic movements indicating decisive action toward a known location; and *Hesitation*, marking periods of deceleration or erratic movement outside target zones that reflect uncertainty or trajectory correction. From these segmented states, we extracted state-specific metrics such as the fraction of time spent in *Search* mode and the average velocity during *Travel*.

All individual features (N=26; see Table S1) were extracted for the whole trial and until target 10 (see ‘Data Quality Checks’ section), and for Part-A and Part-B separately, completing a total of 104 features (26 × 2 × 2). These 104 features were used as input of the Machine Learning pipeline, which includes a Feature Selection step (see Figure 3 and section ‘Machine Learning’ below).

**Fig. 3.**
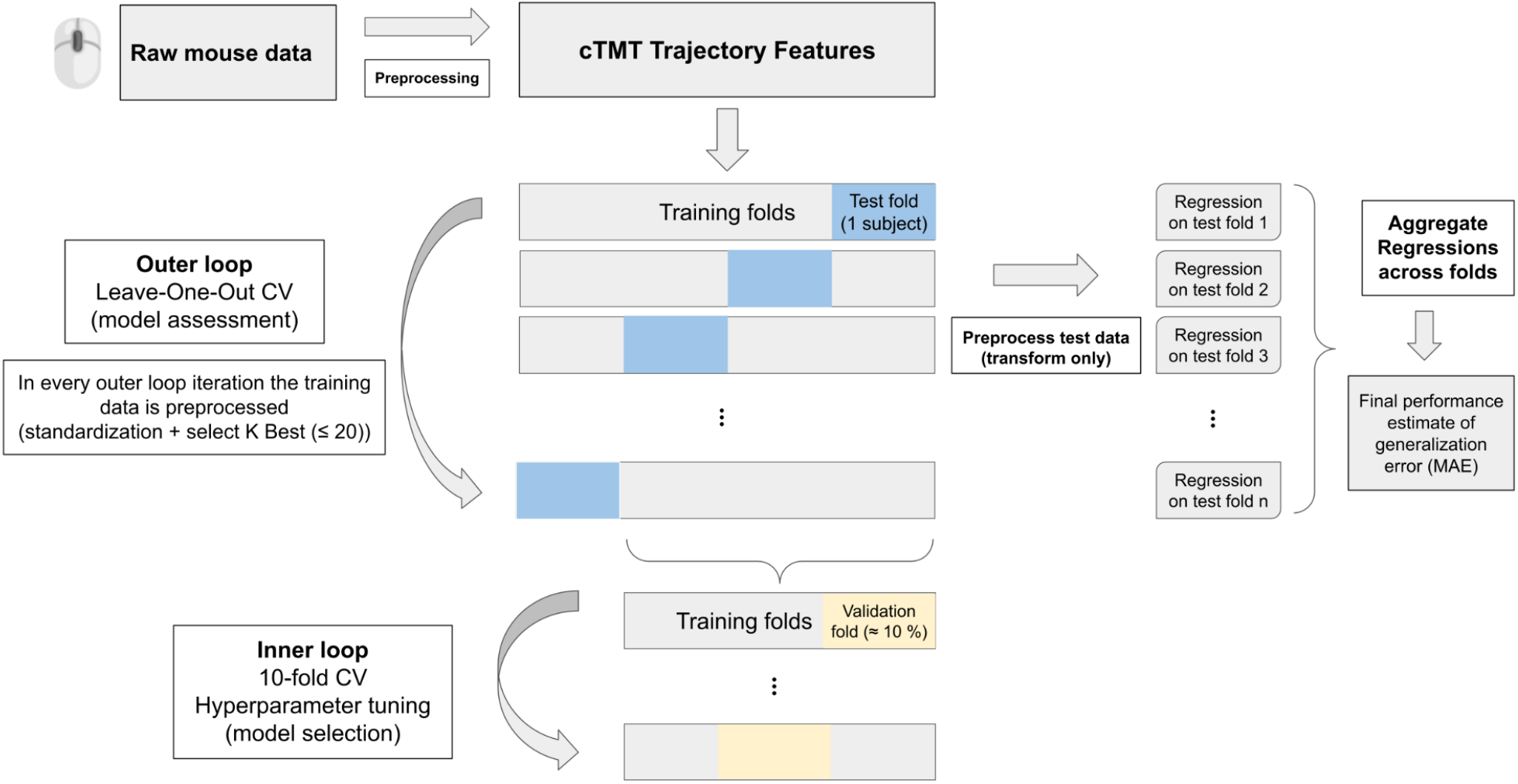
Nested cross-validation framework for regression analysis. The outer loop employs Leave-One-Out Cross-Validation (LOOCV), where each subject is held out once as the test set. Within each outer iteration, the training data is preprocessed through a sequential pipeline: mean imputation of missing values, univariate feature selection (SelectKBest, k ≤ 20, scored by F-regression), and standardization. These transformations are fit exclusively on the training partition and applied to the held-out test subject to prevent data leakage. The inner loop performs 10-fold cross-validation for hyperparameter tuning via grid search, optimizing negative mean absolute error. Predictions from all LOOCV folds are aggregated to compute mean absolute error (MAE) as the final performance estimate.

### F. Target Variables

To train the predictive models, we reduced the high-dimensional output of the validation tasks into single scalar metrics representing core executive functions. Visual working memory was defined as the visual working memory capacity (Cowan’s K) derived from the Change Detection Task, for the 4 (K_4_) and 6 (K_6_) items trials separately, and the averaged (*K*_*mean*_) (Table 1). Inhibition was quantified using the Stop Signal Reaction Time (SSRT) calculated from the Stop-Signal Task (Table 1) and several measures derived from the Go/No-Go task. We present results for the *c*-coefficient and accuracy (Table 1). Finally, we explored the Age as Target Variable, as it is known that it is tightly related with TMT behavior ^32^.

### G. Machine Learning

Following the methodology validated in our previous work ^33^, we implemented a nested cross-validation (CV) framework to obtain unbiased performance estimates across all models (Fig. 3). The outer loop used Leave-One-Out CV (LOOCV) to evaluate generalization performance. In each iteration, one subject was held out as the test fold, and the remaining subjects formed the training set. The preprocessing pipeline consisted of four sequential steps applied within each outer fold to prevent data leakage ^34^: (1) missing value imputation using the training-set mean (SimpleImputer), (2) univariate feature selection via SelectKBest (scikit-learn v1.7.1; ^35^), retaining up to 20 features ranked by their F-scores (using F-regression test), (3) standardization to zero mean and unit variance (StandardScaler), and (4) model fitting.

Within each outer training set, we performed 10-fold CV (inner loop) for hyperparameter tuning and model selection. Hyperparameters were optimized via grid search (GridSearchCV) using the negative mean absolute error for regression, based on the recommendations of Poldrack and collaborators ^36^. After optimal parameters were selected, the pipeline was retrained on the entire outer-training partition and evaluated on the left-out subject. Predictions were aggregated across all LOOCV folds to compute mean absolute error (MAE) as final performance metric. Poldrack et al. (2020) also explicitly caution against using correlation coefficients or R-squared as measures of predictive performance in regression analyses, noting that correlation measures linear association but not predictive accuracy ^36^. Instead, we report the normalized error (MAE / SD) as an effect size metric, which expresses the prediction error relative to the natural variability of each test score and allows comparison across tests with different scales. All experiments used fixed random seeds for reproducibility. This methodology is crucial for obtaining unbiased performance estimates, as it strictly separates the data used for model selection from the data used for performance evaluation ^37^.

We evaluated seven models spanning different families: Linear Regression, Ridge, Lasso, Elastic Net, Support Vector Regression (SVR), Random Forest, and XGBoost. The “no-free-lunch” theorems ^38^ assert that no single algorithm consistently outperforms all others across every problem domain, motivating this multi-model comparison. Complete hyperparameter grids for all models are provided in Table S2 of the Supplementary Materials.

### H. Statistical analysis

The p-value for each model and target variable (see Table 2) was estimated with a permutation test. The test compares the observed MAE with the MAE estimated in N = 1000 permutations of the association between the target variable values and the feature vector for each individual (eq. 1).

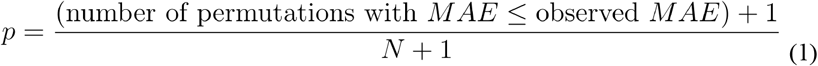

**Table 2.** Each model comprises a different subset of participants who completed both tasks (see sample sizes in Methods). Bold letters indicate the models with lower MAE (and then lower p-value to resolve the draws). See Supplementary Materials for all experiments. Subjects’ Subset refers to the participants that completed the tasks, for instance ‘cTMT ∩ CDT’ indicates the participants that completed both the cTMT and the CDT. The IQR and Standard Deviation were estimated on the observed data, and used for comparison with the MAE. The normalized error was estimated as the ratio between the MAE and the SD. The 95% CIs are bootstrap-based (N = 10,000). The p-value was estimated with a permutation test (N=1000, see Methods).

| Target | Subjects' Subset | Model | MAE | 95% CI (MAE) | SD<sub>Target</sub> | Norm. error (MAE/SD) | IQR<sub>Target</sub> | p-value |
| --- | --- | --- | --- | --- | --- | --- | --- | --- |
| **age** | **cTMT** | **SVR** | **6.78** | **[6.10, 7.53]** | **10.06** | **0.67** | **11.0** | **<0.001** |
| age | cTMT | Ridge | 7.08 | [6.46, 7.74] | 10.06 | 0.7 | 11.0 | <0.001 |
| age | cTMT | ElasticNet | 7.08 | [6.47, 7.74] | 10.06 | 0.7 | 11.0 | <0.001 |
| age | cTMT | LinearRegression | 7.18 | [6.55, 7.85] | 10.06 | 0.71 | 11.0 | <0.001 |
| age | cTMT | Lasso | 7.19 | [6.58, 7.84] | 10.06 | 0.71 | 11.0 | <0.001 |
| age | cTMT | RandomForestRegressor | 7.33 | [6.71, 7.98] | 10.06 | 0.73 | 11.0 | <0.001 |
| age | cTMT | XGBRegressor | 7.61 | [6.91, 8.34] | 10.06 | 0.76 | 11.0 | <0.001 |
| **$K_{mean}$** | **cTMT ∩ CDT** | **RandomForestRegressor** | **0.57** | **[0.51, 0.64]** | **0.72** | **0.79** | **0.94** | **0.011** |
| $K_{mean}$ | cTMT ∩ CDT | Ridge | 0.57 | [0.51, 0.63] | 0.72 | 0.79 | 0.94 | 0.024 |
| $K_{mean}$ | cTMT ∩ CDT | XGBRegressor | 0.59 | [0.53, 0.65] | 0.72 | 0.82 | 0.94 | 0.003 |

**Table 2 (Cont.).** Each model comprises a different subset of participants who completed both tasks (see sample sizes in Methods).
| Target | Subjects’ Subset | Model | MAE | 95% CI (MAE) | SD<sub>Target</sub> | Norm. error (MAE/SD) | IQR<sub>Target</sub> | p-value |
| --- | --- | --- | --- | --- | --- | --- | --- | --- |
| **accuracy** | **cTMT ∩ GNG** | **SVR** | **0.08** | **[0.07, 0.08]** | **0.09** | **0.89** | **0.16** | **0.003** |
| accuracy | cTMT ∩ GNG | RandomForestRegressor | 0.08 | [0.07, 0.08] | 0.09 | 0.89 | 0.16 | 0.004 |
| accuracy | cTMT ∩ GNG | Ridge | 0.08 | [0.07, 0.08] | 0.09 | 0.89 | 0.16 | 0.007 |
| accuracy | cTMT ∩ GNG | ElasticNet | 0.08 | [0.07, 0.08] | 0.09 | 0.89 | 0.16 | 0.012 |
| accuracy | cTMT ∩ GNG | XGBRegressor | 0.08 | [0.07, 0.09] | 0.09 | 0.89 | 0.16 | 0.015 |
| accuracy | cTMT ∩ GNG | Lasso | 0.08 | [0.07, 0.08] | 0.09 | 0.89 | 0.16 | 0.020 |
| accuracy | cTMT ∩ GNG | LinearRegression | 0.08 | [0.07, 0.09] | 0.09 | 0.89 | 0.16 | 0.053 |
| **c-coefficient** | **cTMT ∩ GNG** | **Ridge** | **0.43** | **[0.38, 0.48]** | **0.6** | **0.72** | **0.66** | **<0.001** |
| c-coefficient | cTMT ∩ GNG | Lasso | 0.43 | [0.38, 0.48] | 0.6 | 0.72 | 0.66 | <0.001 |
| c-coefficient | cTMT ∩ GNG | ElasticNet | 0.44 | [0.39, 0.49] | 0.6 | 0.73 | 0.66 | <0.001 |
| c-coefficient | cTMT ∩ GNG | RandomForestRegressor | 0.45 | [0.40, 0.50] | 0.6 | 0.75 | 0.66 | <0.001 |
| c-coefficient | cTMT ∩ GNG | LinearRegression | 0.47 | [0.41, 0.52] | 0.6 | 0.78 | 0.66 | 0.004 |
| c-coefficient | cTMT ∩ GNG | SVR | 0.45 | [0.40, 0.50] | 0.6 | 0.75 | 0.66 | 0.006 |
| c-coefficient | cTMT ∩ GNG | XGBRegressor | 0.5 | [0.45, 0.56] | 0.6 | 0.83 | 0.66 | 0.040 |

### I. Feature importance: SHapley Additive exPlanations (SHAP) approach

The SHapley Additive exPlanations (SHAP) was used to gain deeper understanding of the models’ results ^39^. We computed the SHapley values within each outer LOOCV fold. For each fold, the SHapley values were calculated for the held-out participant, using the training set as background. The chosen model in each fold -with the optimized hyperparameters-was refitted on the training set. The absolute contribution of each feature across all folds were presented as the average SHapley values across folds in which it appeared, since feature selection (SelectKBest) could lead to different subsets of features for different folds. This procedure also yielded more stability of feature importance values.

## III. Results

### A. Behavior data

Following the quality control and feature extraction procedures described above, the final descriptive statistics for each task are presented in Figure 4. For the Computerized Trail Making Task (cTMT), data from 368 participants met the strict quality control criteria (requiring valid completions for both Part A and Part B). The average time to complete Part A was 21.93 s (SD = 2.10), while the more cognitively demanding Part B required an average of 23.82 s (SD = 1.18). A paired t-test confirmed that the increase in duration for Part B was statistically significant (t(367) = -20.48, p < .001, Cohen’s d = 1.07), confirming the expected executive cost. Regarding the Percentage of Completed trials (PC), participants successfully completed, i.e. pass through at least 10 targets in correct order, an average of 88.92% (SD = 15.36) of Part A trials and 73.55% (SD = 22.80) of Part B trials. A paired t-test revealed that this difference was statistically significant (t(367) = 16.10, p < .001, Cohen’s d = 0.84), indicating that the increased cognitive demands of Part B were accompanied by a lower rate of complete trials.

**Fig. 4.**
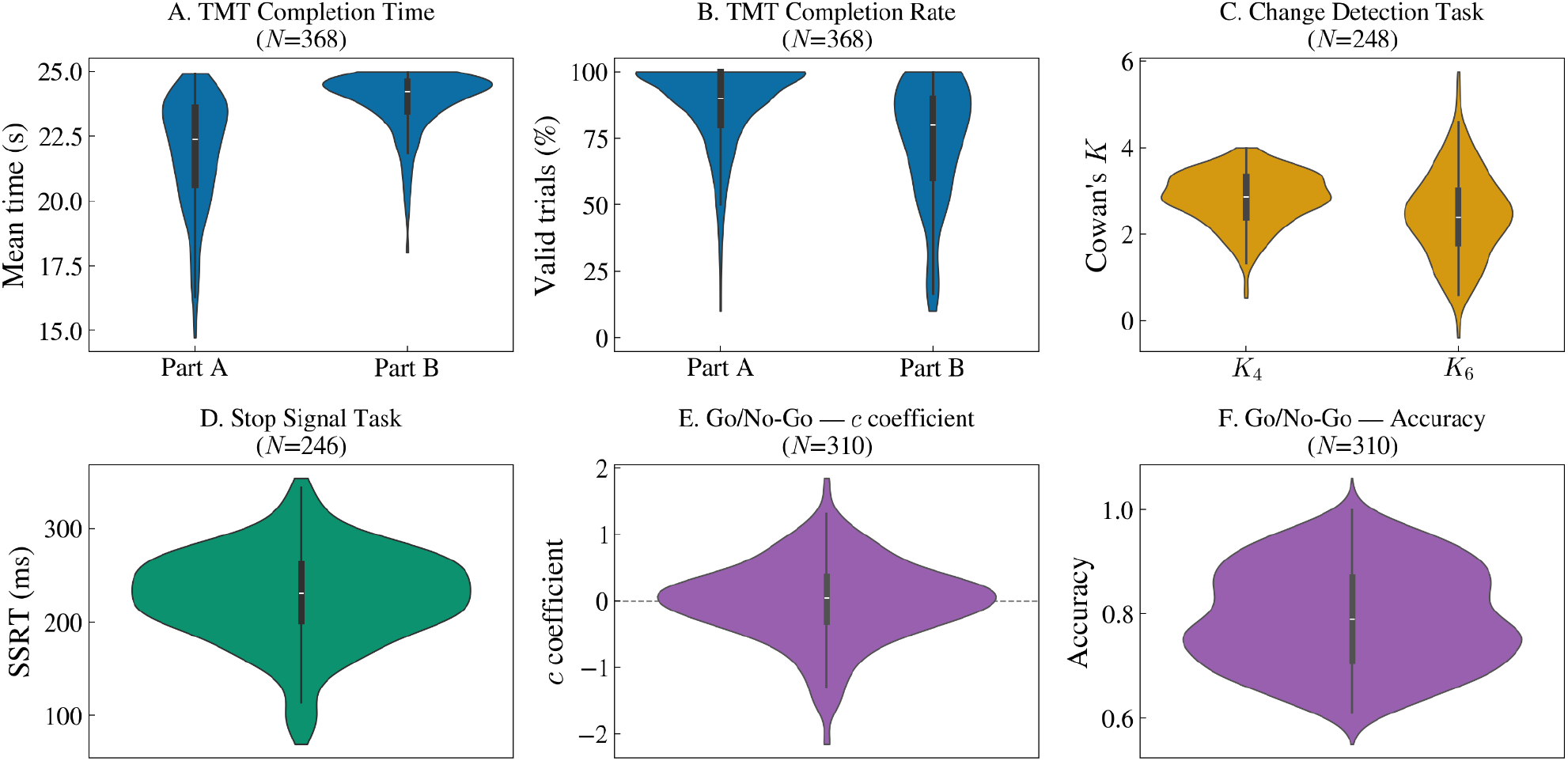
Distribution of performance metrics across cognitive tasks. Violin plots display the kernel density estimate of each metric, with an embedded interquartile box and median marker. **A**. Mean completion time in the cTMT for both parts. **B**. Percentage of valid (completed) trials in the cTMT. **C**. Visual working memory capacity estimated by Cowan’s K for set sizes 4 and 6 in the Change Detection Task (CDT). **D**. Stop-Signal Reaction Time (SSRT), computed via the integration method, as a measure of inhibitory control. **E**. *c*-coefficient from the Go/No-Go task. F. Accuracy in the Go/No-Go task, reflecting the proportion of correct responses across Go and No-Go trials. Colors denote cognitive tasks: blue = cTMT (A, B), amber = CDT (C), green = SST (D), purple = Go/No-Go (E-F). Sample sizes (N) vary across panels due to task-specific quality control exclusion criteria.

Regarding Stop-Signal Task (SST), data from 246 participants met the quality control criteria. The average Stop Signal Reaction Time (SSRT) was 227.7 ms (SD = 49.3), with a mean stop-signal response probability of 0.48. This probability value near 0.5 indicates successful tracking of the response threshold by the adaptive staircase algorithm, ensuring the validity of the SSRT estimation.

In the Go/No-Go Task (GNG), a total of 310 participants were included in the final analysis, after excluding 18 subjects due to low accuracy (<60%). The metrics showed a mean Hit Rate of 0.82 (SD = 0.11) and a False Alarm rate of 0.27 (SD = 0.14), reflecting a moderate level of inhibitory challenge. The mean Reaction Time on Go trials was 333 ms (SD = 26), consistent with standard cognitive control tasks ^40^.

Finally, the analyses for the Change Detection Task (CDT) included 248 participants following the application of quality control procedures. Subjects were excluded if they exhibited a trial omission rate exceeding 20% or a global accuracy lower than 60%. The analysis revealed a mean capacity of 2.83 (SD=0.61) for the low-load condition (set size 4) and 2.45 (SD=1.00) for the high-load condition (set size 6). Global accuracy across the sample was 78%

### B. Prediction results

Different predictive models were implemented in the same training framework for different target variables. Predictive models varied in complexity from Linear Regression to Boosting (XGBRegressor), including regularized regression (Ridge, LASSO, Elastic Net), Support Vector Regression (SVR), and Random Forest Regression. Table 2 summarized the significant results for the selected target variables. Significance was assessed using a permutation test, and MAE comprises the effect size. The MAE should be compared with the dispersion estimates for the target variables, estimated from the sample (SD and IQR).

The cTMT showed strong predictive power for age, as all models yielded significant predictions with p-values < 0.001 (N permutations = 1000) and MAEs one third lower than the dispersion of the target variable. To ensure stable estimates of model performance, we calculated bootstrap 95% confidence intervals (N = 10,000 resamples) for all metrics. Regarding visual working memory, the cTMT measures presented a significant prediction on the *K*_*mean*_. Interestingly, this involves mainly more complex models such as the Random Forest Regressor and XGBRegressor. This suggests that the relation between K and the cTMT behavior could be non-linear and probably involves interactions between the variables. The Go / No-Go was characterized with several target variables, following its vast bibliography, and showed significant effects for all of them. The Accuracy and *c*-coefficient are the ones with stronger effects and the regularized linear models were among the most significant models in both cases, suggesting a linear dependence. The effect sizes were very similar between different models. The Stop-Signal Task, characterized by the SSRT did not present significant results.

### C. Feature Importance Analysis

Feature importance was estimated using mean absolute SHAP values computed for the best-performing model per target variable, selected by lowest permutation test p-value. For age prediction (Fig. 5A), the dominant features were distance-based metrics, specifically complete search distance and complete total distance, present in both Part A and Part B, alongside speed metrics and inter-target time. The relevance of trajectory length is consistent with the concept of “circuitousness” described in the TMT literature ^41^, suggesting that spatial efficiency of cursor paths carries strong individual information about age. The presence of Part A features at the top of the ranking indicates that basic visuomotor execution, independent of alphanumeric switching demands, is particularly informative, consistent with the known sensitivity of motor behavior to lifespan changes ^10,32^.

**Fig. 5.**
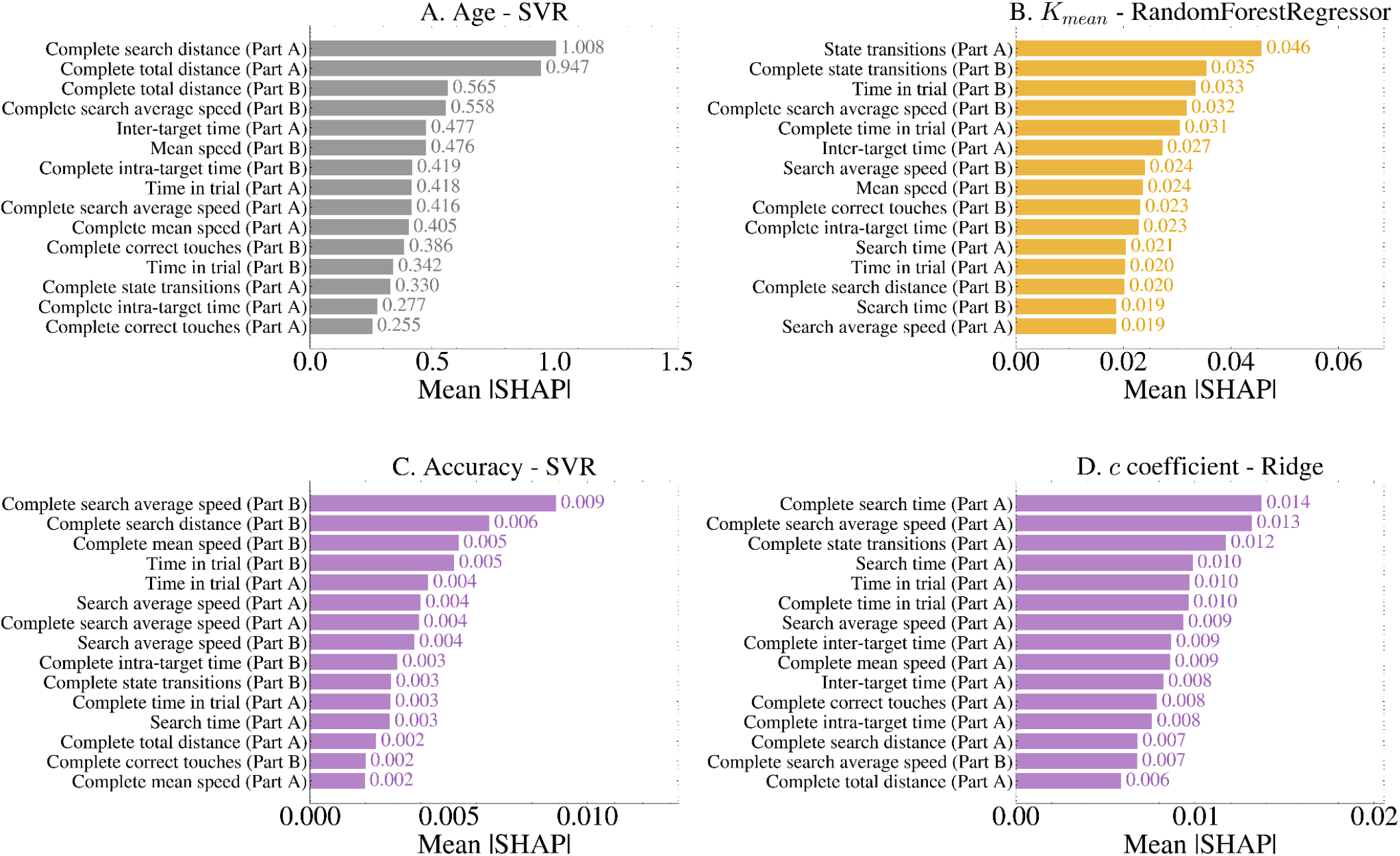
Horizontal bars show mean |SHAP| values across LOO folds for the top 15 features per model. **A**. SVR model prediction on Age; shown in gray. **B**. XGBRegressor model prediction on *K*_*mean*_ (CDT); shown in amber. **C-D**. SVR regression model prediction on Accuracy (C) and Ridge model prediction on *c*-coefficient (D) (Go/No-Go); shown in purple. All models used the feature set from cTMT. Importance scores were averaged across folds, accounting for varying feature subsets due to feature selection (*see Methods*).

For visual working memory capacity (*K*_*mean*_) prediction (Fig. 5B), state transitions in Part A and complete state transitions in Part B emerged as the top predictors with equal importance (0.046 and 0.035 respectively), followed by temporal and speed-based features distributed across both parts. Remarkably, there are several Search attributes between the top 15, mostly related to Part B but with contributions of Part A: Complete search average seed (Part A and B), Search average speed (Part B), Search time (Part A and B), and Complete search distance (Part B). This is in line with it being the VWM most demanding phase. Unlike age prediction, where Part A features dominated, the more balanced contribution of Part A and Part B features here suggests that working memory capacity modulates cursor dynamics across both the visuomotor and switching components of the task. This suggests that *K*_*mean*_ is not just related with functions like task switching (Part B), but also defines the baseline efficiency of visual scanning and trajectory planning throughout the entire test.

Regarding the prediction of the Go/No-Go *c*-coefficient, the SHAP values showed an almost absolute predominance of cTMT-A derived metrics, which accounted for 14 of the 15 variables in the feature importance ranking (Fig. 5D). The most influential variables were “Complete search time (Part A)”, “Complete search average speed (Part A)” and “Complete state transitions (Part A)”, indicating that performance during the simple sequencing task of cTMT-A constitutes the most informative functional domain for predicting response bias.

In the case of the prediction of the Go/No-Go accuracy, a predominance of cTMT-B-derived metrics appeared in the top positions of the feature importance ranking (Fig. 5C). The most influential variables seemed to be “Complete search average speed (Part B)”, “Complete search distance (Part B)” and “Complete mean speed (Part B)”, suggesting that execution efficiency during the cTMT-B alternation task constitutes the most informative functional domain for predicting inhibitory performance. Part A metrics were predominantly ranked in the middle/bottom positions, indicating a lesser contribution of simple sequencing components to the prediction. The overall range of SHAP values was narrow and low in magnitude, reflecting a distributed prediction pattern across multiple features without marked reliance on any single predictor. A similar pattern is observed if the SHAP values were estimated from the Ridge model, which is also significant and provides a more direct comparison with the results for the *c*-coefficient (Supplementary Figure 1).

Considered alongside the *c*-coefficient model, where cTMT-A metrics dominated the importance ranking, the results suggest a differential pattern in which perceptual-motor processes captured by Part A appear more closely linked to response bias, whereas executive processes assessed by Part B are more strongly associated with overall inhibitory performance. This dissociation aligns with evidence that the two parts of the cTMT capture differentiated cognitive processes ^11^.

## IV. Discussion

This study aimed to explore the potential of the computerized Trail Making Test (cTMT) in identifying individual differences in core executive functions by extracting high-resolution features from cursor trajectories. Our results show that cursor dynamics, particularly those related to motor fluency and state transitions, provide a rich behavioral signal for predicting these cognitive capacities, extending the information captured by traditional global completion time metrics. To summarize, the key findings include: (1) a strong relationship between age and cTMT features, especially those reflecting ‘circuitousness’; (2) successful prediction of visual working memory (VWM) capacity (Cowan’s K) using nonlinear models, indicating that VWM acts as a pervasive constraint on search efficiency, even under low executive load conditions (Part A); and (3) a differential prediction pattern for Go/No-Go outcomes, where Part B metrics best predicted inhibitory accuracy while Part A metrics best predicted response bias (*c*-coefficient).

### A. Theoretical implications: cTMT as a continuous cognitive probe

Traditionally, the TMT is viewed as a measure of cognitive flexibility and task switching, with Part B primarily indexing the cost of shifting between alphanumeric sets. However, our feature importance analysis reveals that the “motor” component (Part A) is more cognitively loaded than previously assumed. Notably, state transitions in Part A emerged as the strongest predictor of *K*_*mean*_, matching its Part B counterpart, while additional Part A features such as inter-target time and search average speed contributed moderately. This supports the idea that the TMT is essentially a visual search and trajectory planning task where the demand for goal-directed monitoring is constant ^11^.

Interestingly, the most robust predictions for *K*_*mean*_ were achieved using non-linear models, such as XGBoostRegressor or Random Forest. This suggests that VWM capacity influences cTMT performance through multifaceted non-linear interactions between motor planning and visual search rather than through a direct linear relationship. The importance of the Search attributes is in line with it being the VWM most demanding phase, complementing previous results from eye movements<u>42</u>. Moreover, the prominence of state transitions indicates that the fluency of goal-directed navigation is highly sensitive to memory capacity. This finding aligns with the “processing speed theory” ^43^, but it adds a spatial dimension.

Regarding inhibitory control, the differential predictive pattern observed between overall accuracy and response bias (*c*-coefficient) highlights the multidimensionality of both the Go/No-Go task and the cTMT. While inhibitory accuracy was best predicted by Part B metrics (such as search speed and distance during the alternation task), response bias was almost exclusively associated with Part A metrics (e.g., search time and state transitions). This dissociation suggests that the basic visuomotor sequencing and search efficiency captured by Part A are tightly linked to an individual’s baseline tendency to respond (response bias). Conversely, the ability to successfully inhibit a prepotent response under cognitive challenge (accuracy) relies heavily on the additional cognitive flexibility and switching resources indexed by Part B. Consequently, precise trajectory features allow for the disentanglement of these overlapping, yet distinct, inhibitory processes.

Interestingly, our predictive models did not yield significant results for the Stop-Signal Reaction Time (SSRT). This null finding may suggest that reactive inhibition ^24^ does not strongly share the same kinematic signatures as the continuous visual search and proactive motor planning required in the TMT. While previous studies have successfully captured subtle spatial and temporal adjustments in proactive control paradigms like the Go/No-Go task, the dynamic, trial-by-trial adjustments inherent to the SST’s adaptive staircase algorithm might introduce variability that is harder to map onto the comparatively global, state-based trajectory features we extracted. This result aligns with recent literature emphasizing that reactive and proactive inhibition, though related, rely on distinct temporal and neural mechanisms ^44^, suggesting the need for different trajectory features to capture reactive control.

Finally, the prediction results of the cTMT behavior with Age not only obtained the lower p-values but also, all models achieve significant results. This is in line with previous results on pen-and-paper TMT ^32^ and, with the strong association of the TMT performance with several cognitive functions ^41^, it suggests a relation of the extracted attributes with brain aging. This establishes cTMT hand features and the pipeline proposed here as a potential biomarker for aging and other related neurodegenerative diseases ^33^. Biomarkers of aging derived from techniques such as fMRI or EEG reach comparable performance levels ^45–47^, but require in-place evaluations, more expensive devices, and more complex logistics. Systematic reviews emphasize that prediction accuracy depends heavily on modality, sample size, age range, and model choice ^48,49^.

Among the top-performing models, the success of linear regressions suggests that age-related decline manifests progressively and proportionally in cursor dynamics. As shown by our SHAP analysis, the features with the highest impact are related to distance-based metrics (e.g., complete search distance and total distance) across both A and B parts. The prominent role of trajectory length supports the ‘circuitousness’ phenomenon, where older age is characterized by decreased spatial efficiency, longer exploratory movements, and increased motor noise during goal-directed navigation, irrespective of the cognitive switching load.

### B. Limitations and future directions

The online approach enables us to access larger samples of individuals than in the lab, in more ecologically valid scenarios. Moreover, a thoughtful machine learning pipeline will assure robust results. Nevertheless, there are some limitations that should be considered. First, variability in hardware and browser performance introduced non-uniform sampling rates of mouse trajectories. In online settings, browsers poll mouse positions at lower rates than the operating system, and network latency further compounds this discontinuity ^50^. Linear interpolation mitigated this issue following standard practice ^4^, though future work would benefit from more standardized mouse-tracking sampling procedures, as sampling rates remain highly variable and underreported across studies ^51^. Second, our sample consisted mostly of highly educated young to middle-aged adults, which limits generalizability to clinical populations or older cohorts at high risk for neurodegeneration. It is worth noting that this distribution also precluded the analysis of the influence of cognitive reserve associated with education, given the overrepresentation of higher educational levels in the sample.

Future studies should expand the variability of the sample, and also explore the test-retest reliability of these features and evaluate their sensitivity in longitudinal designs. Additionally, integrating these metrics with other data streams, such as keystroke dynamics or multimodal ecological assessments (incorporating eye-movements or speech), could improve the predictive accuracy of digital cognitive models. In particular, eye movements in particular could provide a more direct measure of the phases identified in the trajectory segmentation (exploring, monitoring, planning), and also behaviors directly related to inhibition or working memory ^42,52–54^. Recent developments have enabled online eye movement recording, though further improvements in precision are needed for reliable measurements in the TMT ^55–59^.

## V. Conclusions

By moving beyond completion time, we have shown that the online cTMT can serve as a powerful, portable tool for evaluating the nuances of executive control and can be also useful for tracking age-related cognitive decline. Our ability to predict specific capacities, such as working memory, distinct inhibitory processes, and age based on high-resolution cursor trajectories highlights the rich signal embedded in continuous motor behavior. Ultimately, leveraging this complexity in digital environments holds significant potential for the future of computational psychiatry, scalable remote assessments, and personalized neuropsychology.

## Acknowledgements

We thank Pablo Laciana and Lara Gauder for their contribution to the development of the DataPruebas platform, as well as Gustavo Verón, Tomás D’Amelio, and Agustín Petroni for their insightful discussions.

Claude (Anthropic) and ChatGPT (OpenAI) were used to assist with manuscript preparation, particularly for improving English language and clarity. All scientific content, analysis, and interpretation remain the sole responsibility of the authors.

## Code and data availability

The datasets generated and analyzed during the current study are available in the Open Science Framework (OSF) repository (https://osf.io/9rnp4/). The code used for the experimental tasks is available on GitHub at https://github.com/NeuroLIAA/Neuro_psych_tasks, and the machine learning analysis pipeline can be accessed at https://github.com/gianluca-capelo/datapruebas_analysis. Additionally, raw cursor time-series from the cTMT were processed using Neurotask, a custom-built Python library designed for high-resolution behavioral analysis, which is publicly available at https://github.com/NeuroLIAA/neurotask.

## Author’s contributions

According to CRediT categories, GEJ and JEK contributes to the Conceptualization; GEJ and GC to the Data curation, and Software; GEJ, GC, and JEK to the Formal analysis, Investigation, Methodology, Validation, Visualization, and Writing – review & editing. JEK to the Funding acquisition, Project administration, Resources, and Supervision; and GEJ to Writing – original draft.

## Abbreviations

TMT: Trail-Making Test
cTMT: computerized version of the TMT
CDT: Change Detection Task
GNG: Go/No-Go
SST: Stop-Signal Tasks
VWM: Visual Working Memory
AUC: Area Under the Curve (feature extraction)
SSRT: Stop Signal Reaction Time
CV: Cross-Validation (CV)
LOOCV: Leave-One-Out CV
MAE: Mean Absolute Error
SVR: Support Vector Regression
SHAP: SHapley Additive exPlanations
SD: Standard Deviation

