## Supplementary Table 1 for "Beyond completion time: predicting individual differences in executive functions using cursor trajectory features in an online Trail Making Test"

**Table S1.** Hand features extracted from the mouse movement time-series. The same variables were computed separately for Part A, Part B, and for the B/A ratio, resulting in three variants of each feature.

| Feature | Description |
| --- | --- |
| Time in trial | Total duration of the trial. |
| Correct touches | Number of correct target touches |
| Wrong target touches | Count of cursor touches on incorrect targets (not the next expected in the sequence) |
| Area difference from ideal | Difference between the area of the actual path and the ideal path. |
| Distance difference from ideal | Difference between the traveled path length and the length of the ideal straight line connecting to the next target. |
| Inter-target time | Sum of time elapsed between successive target touches. |
| Intra-target time | Sum of time spent within the area of a target before leaving it. |
| Mean speed | Average cursor speed. |
| SD speed | Standard deviation of the cursor speed. |
| Peak speed | Maximum cursor speed observed. |
| Number of crosses | Number of times the cursor trajectory crosses itself, excluding intersections between temporally adjacent segments. |
| State transitions | Total number of transitions for one state (hesitation, search or travel) to another |
| Total distance | Total distance traveled by the cursor. |
| Hesitations | Count of distinct hesitation periods detected in the trial. |
| Hesitation max duration | Longest hesitation duration during the trial |
| Hesitation average duration | Mean duration of individual hesitation periods detected in the trial. |
| Hesitation ratio | Proportion of time in hesitation relative to total travel time: hesitation_time / (travel_time + hesitation_time). |
| Hesitation time | Total time spent in the hesitation state during the trial. |
| Hesitation average speed | Average hesitation speed. |
| Hesitation distance | Total distance traveled by the cursor in hesitation |
| Search average speed | Average cursor speed during search movements. |
| Search distance | Total distance traveled during search movements. |
| Search time | Total time spent in the search state. |
| Travel average speed | Average cursor speed during travel segments. |
| Travel distance | Total distance traveled during travel segments. |
| Travel time | Total time spent during travel segments. |
