## Supplementary Table 2 for "Beyond completion time: predicting individual differences in executive functions using cursor trajectory features in an online Trail Making Test"

**Table S2.** Regression hyperparameters.

| Name | Hyperparameter | Values |
|---|---|---|
| **Random Forest Regressor** | n_estimators | 100, 200, 500 |
| | max_depth | None, 8, 16 |
| | min_samples_leaf | 2, 5, 10 |
| | max_features | sqrt, log2 |
| **Support Vector Machine (SVR)** | C | 0.01, 0.1, 1, 10, 100, 1000 |
| | epsilon | 0.01, 0.1, 0.5, 1.0 |
| | gamma (for rbf kernel) | scale, auto |
| **Linear Regression** | No tunable hyperparameters | - |
| **Ridge Regression** | alpha | 0.0001, 0.001, 0.01, 0.1, 1.0, 5, 10, 100, 1000, 10000 |
| **Lasso Regression** | alpha | 0.0001, 0.001, 0.01, 0.1, 1.0, 5, 10.0 |
| **XGBoost Regressor** | n_estimators | 100, 200 |
| | max_depth | 3, 6 |
| | learning_rate | 0.1, 0.3 |
| | subsample | 0.8, 1.0 |
| | colsample_bytree | 0.8, 1.0 |
| **Elastic Net** | alpha | 0.0001, 0.001, 0.01, 0.1, 1, 10, 100 |
| | l1_ratio | 0.05, 0.2, 0.5, 0.8, 0.95 |
