## Supplementary Figure 1 for "Beyond completion time: predicting individual differences in executive functions using cursor trajectory features in an online Trail Making Test"

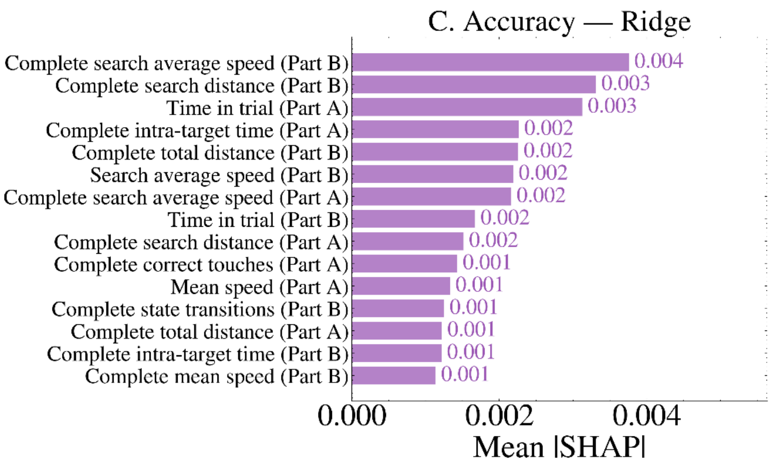


**Figure S1.** SHAP features importance for cTMT-based prediction models. Horizontal bars show mean |SHAP| values across LOO folds for the top 15 features per model. Ridge regression model prediction on Accuracy. Importance scores were averaged across folds, accounting for varying feature subsets due to feature selection (see Methods).
